# Differential gene expression and the importance of regulatory ncRNAs in acidophilic microorganisms

**DOI:** 10.1101/538918

**Authors:** Daniela S. Aliaga Goltsman, Loren Hauser, Mauna Dasari, Brian C. Thomas, Jillian F. Banfield

**Author notes:** Corresponding authors: Daniela S. Aliaga Goltsman, Stanford University, School of Medicine, Department of Infectious Diseases, 3801 Miranda Ave., Bldg. 101, Rm B4-185, Palo Alto, CA 94304, Jillian Banfield, University of California Berkeley, Earth and Planetary Sciences Department Energy Biosciences Building, 2151 Berkeley Way, Berkeley, CA 94720.

## Abstract

Gene expression profiles provide insight into how microorganisms respond to changing environmental conditions. However, few studies have integrated expression profile analyses of both coding genes and non-coding RNAs (ncRNAs) to characterize the functional activity of microbial community members. Here, we defined gene expression profiles from environmental and laboratory-grown acidophilic biofilms using RNASeq. In total, 15.8 million Illumina reads were mapped to the genomes of 26 acidophilic microorganisms and nine viruses reconstructed from the Richmond Mine at Iron Mountain, California. More than 99% of the genome was transcribed in three *Leptospirillum* species, and > 80% in the archaea G-plasma and *Ferroplasma* Type II. High gene expression by G-plasma and the *Leptospirillum* Group II UBA strain correlated with extremely acidic conditions, whereas high transcriptional expression of *Leptospirillum* Group III and *Leptospirillum* Group II 5way-CG strain occurred under higher pH and lower temperature. While expression of CRISPR Cas genes occurs on the sense strand, expression of the CRISPR loci occurs on the antisense strand in the *Leptospirilli*. A novel riboswitch associated with the biosynthetic pathway for the osmolyte ectoine was upregulated when each specific *Leptospirillum* Group II strain was growing under the conditions most favorable for it. Newly described ncRNAs associated with CO dehydrogenase (CODH) suggest regulation of expression of CODH as a CO sensor in mature biofilms in the *Leptospirilli*. Results reveal the ways in which environmental conditions shape transcriptional profiles of organisms growing in acidophilic microbial communities and highlight the significance of ncRNAs in regulating gene expression.

**IMPORTANCE:** Microorganisms play important roles in environmental acidification and in metal-recovery based bioleaching processes. Therefore, characterizing how actively growing microbial communities respond to different environments is key to understanding their role in those processes. Microorganisms express their genes, both coding and non-coding, differently depending on environmental factors, thus evaluating community expression profiles inform about the ecology of actively growing microorganisms. Here we used community transcriptomic analyses to characterize gene expression profiles from biofilm communities growing under extremely acidic conditions. Results expand our knowledge of how acidophilic microorganisms respond to changes in their environment and provide insight into possible gene regulation mechanisms.

## INTRODUCTION

Extremely acidic environments are usually dominated by relatively few taxa making them good model systems for ecology and physiology studies. Because of the roles acidophilic microorganisms play in environmental acidification and in metal-recovery based bioleaching processes, gene expression studies are key to understanding the physiology and ecology of microorganisms in acidic environments. The Richmond Mine at Iron Mountain, California, is a well-studied acid mine drainage (AMD) system: for example, deep sequencing of many Richmond Mine biofilms has enabled reconstruction of the genomes of many Bacteria, Archaea, viruses, plasmids, and fungus (e.g. (1–4)). Genome reconstruction analyses, along with high-throughput transcriptomic sequencing and mass spectrometry-based proteomics have the potential to provide information about the community composition and activity in natural ecosystems. Community transcriptomic analyses have been used to describe important metabolic processes in natural communities such as nitrogen metabolism in marine environments (5), the flow of carbon between organisms in photosynthetic microbial mats (6), and associations of microbial communities with the biogeochemistry of permafrost thawing environments (7). In acidic environments, mass spectrometry-based community proteomics measurements of the dominant member of Richmond Mine AMD biofilms indicated that biofilm maturation affects *Leptospirillum* group II protein expression profiles, with the most protein expression variation found as biofilms transition from early and mid-developmental to the most mature growth stage (8). Similarly, microarray-based community transcriptomics indicated that expression profiles of genes involved in biofilm formation, chemotaxis, motility and quorum-sensing in *Leptospirillum ferrooxidans* were up-regulated when living as part of biofilms vs. in the planktonic fraction of the Rio Tinto AMD system (9). Hua *et al.* recovered the genome of *Ferrovum* spp. from an acid mine drainage system and community gene expression profiles indicated active CO_2_ fixation and sulfate reduction pathways (10). In addition, metatranscriptomics studies have highlighted the importance of non-coding RNA (ncRNA) expression in marine systems (11). However, studies of the transcriptional profiles of coding and ncRNAs in acidophilic communities relevant for AMD generation and bioleaching-based metal recovery have not yet been reported.

Many ncRNAs have been identified in Bacteria and Archaea, such as riboswitches, ribozymes, and other regulatory ncRNAs, which play important roles in regulating gene expression (reviewed in (12)). Riboswitches are regions within a messenger RNA (generally located in the 5’ untranslated region - UTR) containing ligand-binding sensors that regulate downstream coding sequences (reviewed in (13)). Other regulatory ncRNAs with known function include the tmRNA, which helps unlock stalled ribosomes, and the 6S ncRNA, which acts as mimic of a promoter (reviewed in (14)). In Eukaryotes, a common mechanism of gene regulation involves transcription of long 5’ UTRs with secondary structures that can influence transcription of downstream genes, or containing short upstream open reading frames (uORFs) that attenuate translation of the downstream protein (reviewed in (15)). In addition, high-throughput transcriptomics analyses have enabled identification of ncRNAs that regulate antibiotic resistance in bacteria via 5’ UTR transcription and uORF attenuation (reviewed in (16)).

Here we report the analyses of transcriptional profiles of non-ribosomal RNA from biofilms collected from the Richmond Mine at Iron Mountain, California, and from laboratory-grown biofilm communities. Gene expression profiles were evaluated for the whole acidophilic community including Bacteria, Archaea, and viruses, and novel ncRNAs were discovered for many community members. The results greatly expand our understanding of the responses of acidophilic organisms to changes in their environment and provide insight into possible gene regulation mechanisms by ncRNAs in the *Leptospirilli*.

(Part of this article was submitted to an online preprint archive (17))

## RESULTS

### Transcriptomics reads map across the whole genome of AMD community members

On average 91.82% ± 4.35% of the transcriptomic reads in total RNA samples and 79.80% ± 9.59% in the rRNA-subtracted samples mapped to rRNA genes from the SSU and LSU Silva databases (Table 1). Despite the low efficiency of the rRNA-depletion protocol used, deep sequencing allowed us to detect transcripts from 37 near-complete and partial genomes of Bacteria, Archaea, Fungi, plasmids and viruses (Supplementary Table S1). Transcriptomic data indicate that *Leptospirillum* group II C75 and UBA genotypes dominate early and mid-developmental stage environmental biofilms characterized by extremely low pH and high temperature, whereas *Leptospirillum* group II 5way-CG is the most abundant genotype in the late developmental-stage environmental biofilm (Figure 1A). *Leptospirillum* group III bacteria dominate bioreactors and the A-drift environmental sample. *Leptospirillum ferrooxidans* and *Acidithiobacillus caldus* were detected at very low abundance, as were other organisms and viruses, some of which represent less than 0.001% of the community transcriptome (Supplementary Table S1). Low Archaeal gene expression and high *Acidithiobacillus* sp. expression have been documented on acid mine drainage biofilms growing at higher pH (18).

**Table 1.** Summary statistics and description of samples sequenced. Samples were collected directly from the Richmond Mine (Env) or from bioreactor-grown biofilms (BR). GS: growth stage; Temp: temperature; Type of sequencing: GAII, Illumina GAIIx platform; HiSeq, Illumina HiSeq 2500 platform.

| Sample ID | Date collected | Env Type | GS | Location | Temp | pH | Type seq. | Total RNA |  | rRNA-subtracted |  | Non-rRNA Reads (M) |
| --- | --- | --- | --- | --- | --- | --- | --- | --- | --- | --- | --- | --- |
|  |  |  |  |  |  |  |  | No. reads (M) | % rRNA | No. Reads (M) | % rRNA |  |
| R1_GS0 | 3/30/10 | BR | 0 | Outflow | 37 | NA* | GAI | 2.56 | 90.55 | 4.27 | 84.85 | 0.89 |
| R1_GS05 | 2/19/10 | BR | 0.5 | Outflow | 37 | 1.65 | GAI | 4.86 | 88.34 | - | - | 0.57 |
| R2_GS05 | 7/20/09 | BR | 0.5 | A drift | 37 | NA* | GAI | 4.79 | 84.97 | - | - | 0.72 |
| A-drift | 9/17/10 | Env | 0 | A drift | 40 | 1.27 | GAI | 3.41 | 86.00 | - | - | 0.48 |
| C75_GS1 | 9/17/10 | Env | 1 | C drift | 46 | 0.86 | GAI | 4.66 | 90.58 | 4.59 | 77.17 | 1.49 |
| AB10_GS0 | 11/2/10 | Env | 0 | AB drift | 39 | 0.8 | HiSeq | 27.87 | 96.22 | 53.15 | 82.09 | 10.57 |
| AB10_GS1 | 11/2/10 | Env | 1 | AB drift | 39 | 0.8 | HiSeq | 30.24 | 96.46 | 58.42 | 81.82 | 11.69 |
| C10_GS0 | 11/2/10 | Env | 0.5 | C drift | 42 | 0.8 | HiSeq | 30.91 | 94.12 | - | - | 1.60 |
| C10_GS05 | 11/2/10 | Env | 0 | C drift | 42 | 0.8 | HiSeq | 27.12 | 96.16 | 51.88 | 85.39 | 8.77 |
| C10_GS1 | 11/2/10 | Env | 1 | C drift | 42 | 0.8 | HiSeq | 28.60 | 95.49 | 50.65 | 85.66 | 8.55 |
| G2E1_GS2 | 6/9/09 | Env | 2 | 4-way | 39 | 0.7 | HiSeq | 26.68 | 85.73 | 45.06 | 68.76 | 17.88 |
| R3_GS0 | 9/28/10 | BR | 0 | A drift | 37 | 1.31 | HiSeq | 31.56 | 95.00 | - | - | 1.58 |
| R3_GS1 | 10/6/10 | BR | 1 | A drift | 37 | 1.74 | HiSeq | 30.49 | 94.08 | 67.98 | 75.10 | 18.73 |

\* NA indicates that no pH measurements were available for these samples.

**Figure 1.**
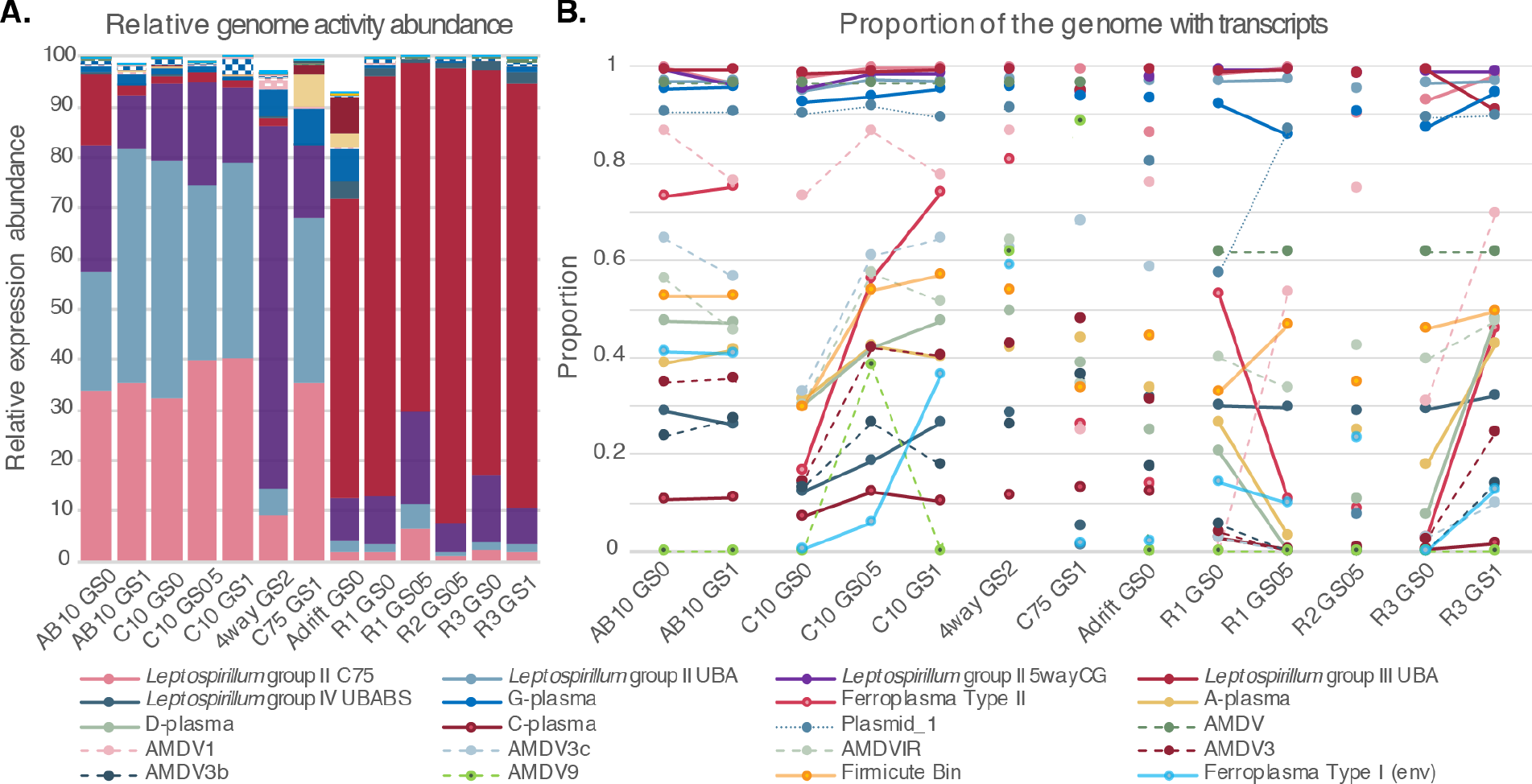
Differential gene and genome expression correlates with environmental conditions in AMD biofilms. A) Relative expression abundance of AMD genomes. Read counts were normalized by the genome length and sample size. B) Proportion of AMD genomes covered by transcriptomic reads. The top 20 most abundant taxa and viruses are shown.

Transcriptomic reads spanned > 95% of the genome of *Leptospirillum* group II and group III, the G-plasma archaeon and one viral genome (Figure 1B). In addition, up to 80% of the genome of *Ferroplasma* Type II, and over 60% of the genomes of *Ferroplasma* Type I and three viruses were identified as transcribed in some samples (Figure 1B). Likely, higher sequencing coverage and/or higher rRNA-removal efficiency would support detecting whole genome transcription in low-abundance community members. The results suggest that for dominant and low-activity organisms and viruses the whole genome is transcribed at some level. This trend has been observed previously in transcriptomics analyses of *Bacillus anthracis* isolates (19).

### Gene expression profiles from AMD taxa correlate with the environment pH

Mapping reads to the predicted genes of AMD organisms, plasmids, and viruses were assembled to yield over 26,300 transcripts (Supplementary Materials – R data). Differential expression analysis identified >4,800 significantly differentially expressed genes with a false discovery rate of 0.05 when evaluating environmental vs. bioreactor-grown biofilms (Supplementary Figure S1A). Non-metric multidimensional scaling (NMDS) ordination on the ~4,800 differentially expressed genes indicates two main axes responsible for most of the sample variation (Figure 2, and Supplementary Figure S1B). The NMDS samples plot shows that axis 1 separates primarily based on environment type, while axis 2 separates based on pH (Figure 2A). Gene expression profiles from *Leptospirillum* group II UBA and C75 genotypes appear correlated with negative NMDS2 scores (low pH environments), while expression profiles from *Leptospirillum* group II 5way-CG genotype and group III appear correlated with positive NMDS2 scores (higher pH environments) (Figure 2B-D). Gene expression profiles from Archaea and viruses generally correlated with positive NMDS1 scores, that is, from environmental rather than bioreactor-grown biofilms. Overall, the NMDS genes plots show a gradient of genes from bacteria living at very low pH and high temperature (Figure 2B, bottom right quadrant), transitioning towards genes from Archaea, viruses and other low-abundance organisms (Figure 2C-D), and ending on genes from bacteria living in bioreactors at higher pH and lower temperature (Figure 2B, top left quadrant). Results indicate that environmental gradients directly affect the transcriptional profiles of the community members living along them. Gradients of pH, temperature, and dissolved oxygen, among others, have been suggested as factors influencing community assembly in acidic environments (20, 21).

**Figure 2.**
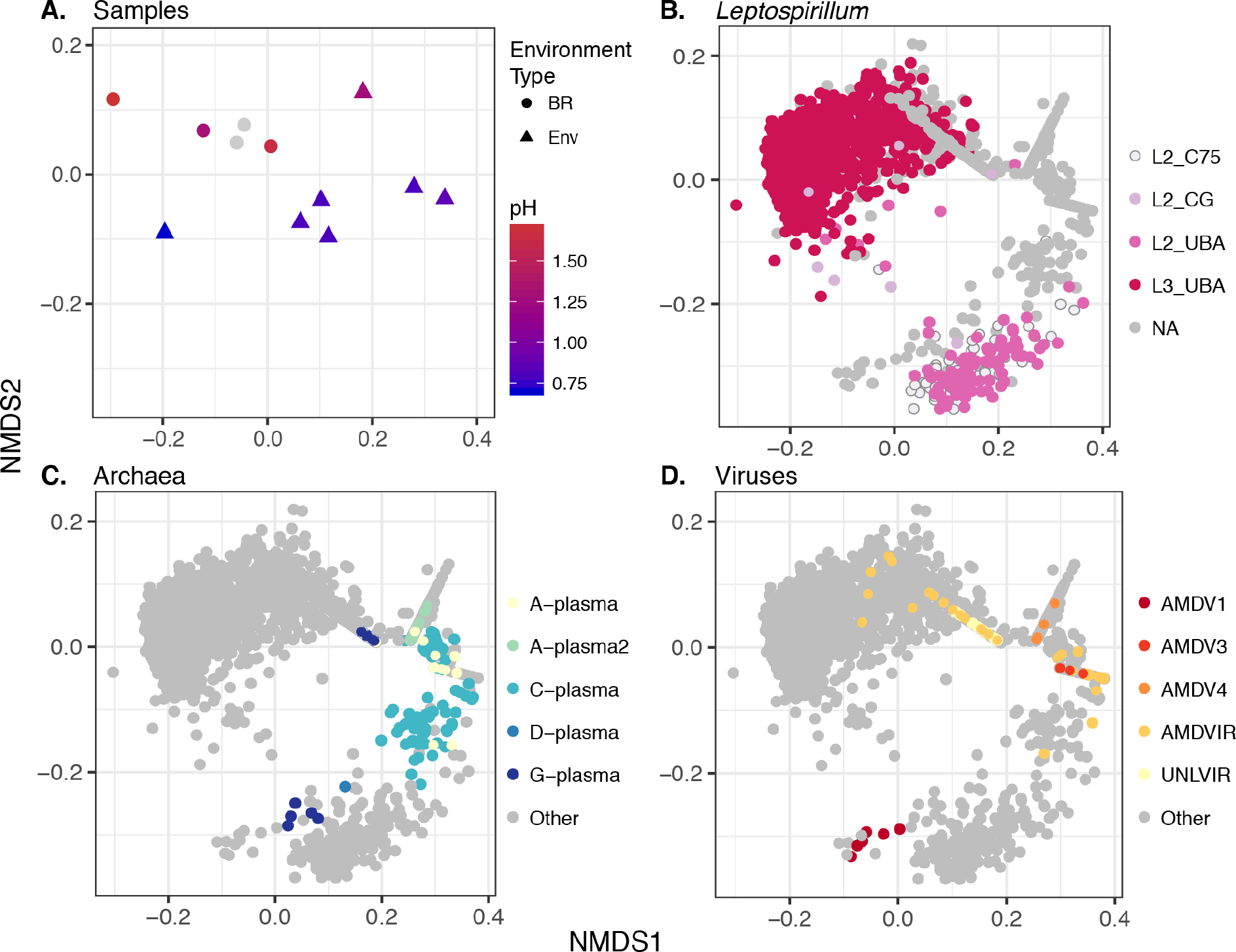
Non-metric multidimensional scaling (NMDS) ordination on Bray-Curtis distances of differentially expressed genes in AMD communities. A) NMDS samples plot color coded by pH and environment type. B-D) NMDS genes plot colored by: B) the top most abundant *Leptospirillum* species and strains; C) the top most abundant Archaeal species, and D) viral genomes. NMDS genes plots indicate a gradient of points going from the lower right quadrant to the top right quadrant, which then goes to the top left quadrant (horseshoe).

Environment-specific transcriptional profiles are also evident from hierarchical clustering of transcribed genes with a log2-fold change >5, which separates samples into environmental and bioreactor biofilms, and environmental samples appear to also cluster by developmental stage (Supplementary Figures S1C and S1D). For example, *Leptospirillum* group II 5way-CG and *Leptospirillum* group III genes are overrepresented in the bioreactors and in the A-drift environmental sample (Supplementary Figures S1D). The A-drift biofilm was collected from a very oxidized pool, at higher pH and lower temperature than other environmental samples (see Table 1), conditions that appear to favor growth of these organisms in the laboratory bioreactors. Genes from *Leptospirillum* group II UBA are over-represented in early to mid-growth-stage environmental biofilms relative to bioreactor-grown biofilms (Supplementary Figure S1D). This genotypic group is also well represented in biofilms collected from the C-drift locations, many of which have reported very low pH and high temperature environments (22–24). Our results support previous proteomic-based studies that suggested distinct ecological adaptation of two *Leptospirillum* group II strains (22). *Leptospirillum* group II-associated AMDV1 phage genes, as well as genes from the AMDV3 virus are overrepresented in the environmental biofilms, while unassigned-viral genes (AMDVIR, likely *Leptospirillum*-associated phage) are overrepresented in bioreactors (Supplementary Figure S1C and Figure 2D). Results indicate that phage/virus activity correlates with their host gene expression.

When looking at the transcriptional profiles of the most abundant community members, hierarchical clustering of expressed genes indicates that genes involved in energy production and conversion, carbon fixation, fatty acid metabolism, transcription and translation factors, and ribosomal proteins from *Leptospirillum* group II UBA and C75 are highly expressed in early to mid-growth-stage environmental biofilms (Figure 3A-B, 3E). These findings suggest rapid growth during early and mid-successional stages. Genes generally overrepresented in bioreactor samples include those involved in amino acid and cofactor metabolism, carbohydrate and lipid metabolism, DNA repair and recombination, lipopolysaccharide metabolism, nucleic acid metabolism, signal transduction, tRNA synthetases and transport genes from *Leptospirillum* group II 5way-CG and group III (Figure 3C-D and 3E). Bioreactors usually have higher pH and lower temperature than most environmental biofilms from the Richmond Mine (Table 1). These conditions might slow down the activity of *Leptospirillum* group II UBA and C75, and of G-plasma, while enriching for growth and transcriptional activity of *Leptospirillum* group III and group II 5way-CG.

**Figure 3.**
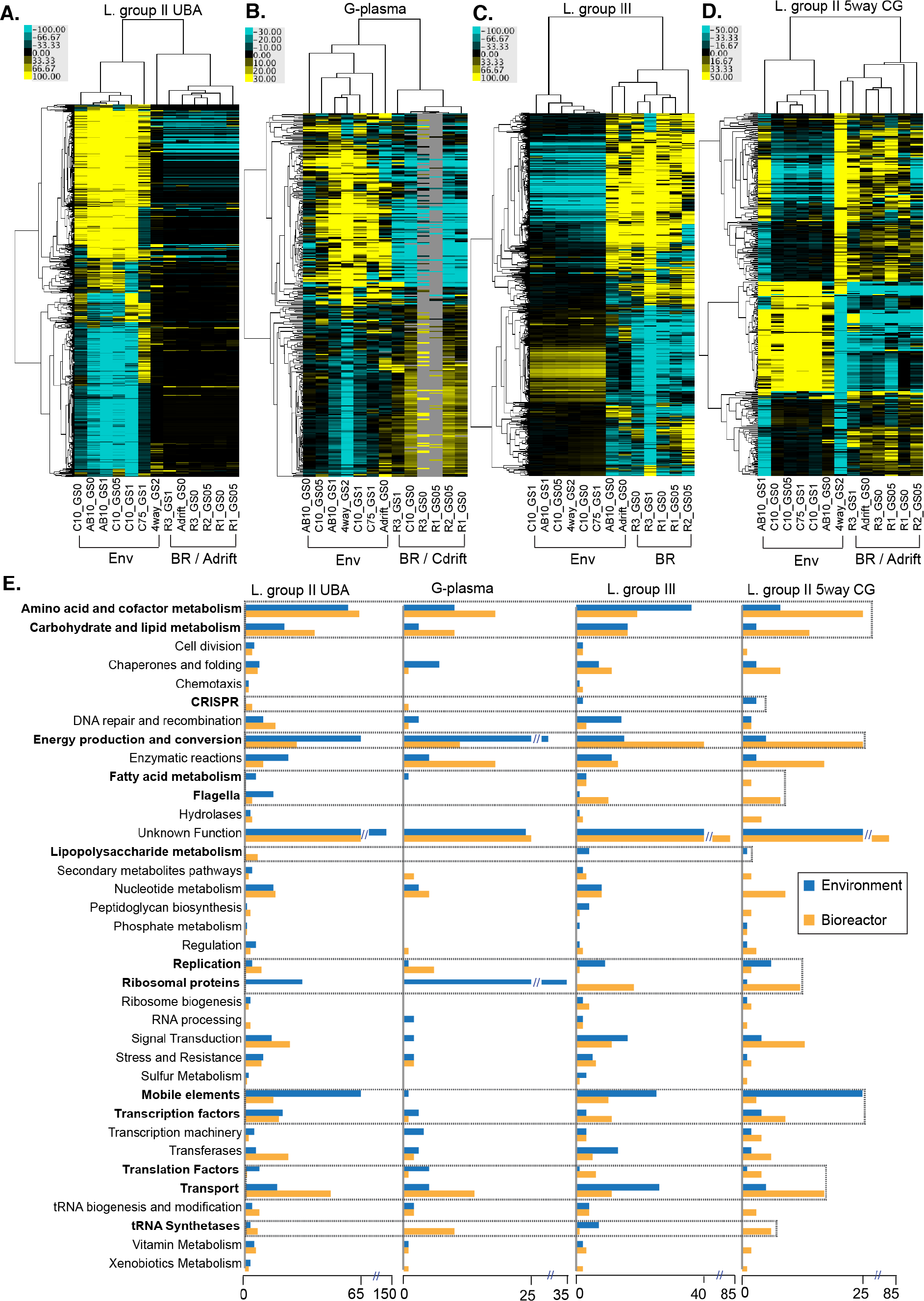
(previous page) Environmental conditions distinguish expression of groups of genes and functional categories in AMD systems. A-D) Heatmaps of genes with log2-fold change > 5 in: A) *Leptospirillum* group II UBA; B) G-plasma; C) *Leptospirillum* group III; and D) *Leptospirillum* group II 5way CG. Yellow: overrepresented, blue: underrepresented. E) Functional categories differ in communities growing in the natural environment vs. laboratory bioreactors. Categories enriched in environmental samples are shown in dark blue, while categories enriched in bioreactor samples are shown in orange. Functional categories in bold within black boxes represent the categories in which the most variation between conditions was observed.

### Regulatory motifs, mobile elements and expression of ncRNAs of known function

Inspection of the upstream regions of transcribed coding genes, tRNAs and non-coding RNAs predicted 207 RpoD promoters in the *Leptospirillum* group II UBA genotype (Supplementary Table S2): 11.5% of them were associated to coding genes or operons transcribed in at least 10 of the 13 biofilm samples. Promoter length varies between 8 and 33 bp and a sequence logo (25) representation of the overall promoter structure indicates that most promoters contain a −10 and −35 motif (Figure 4A).

**Figure 4.**
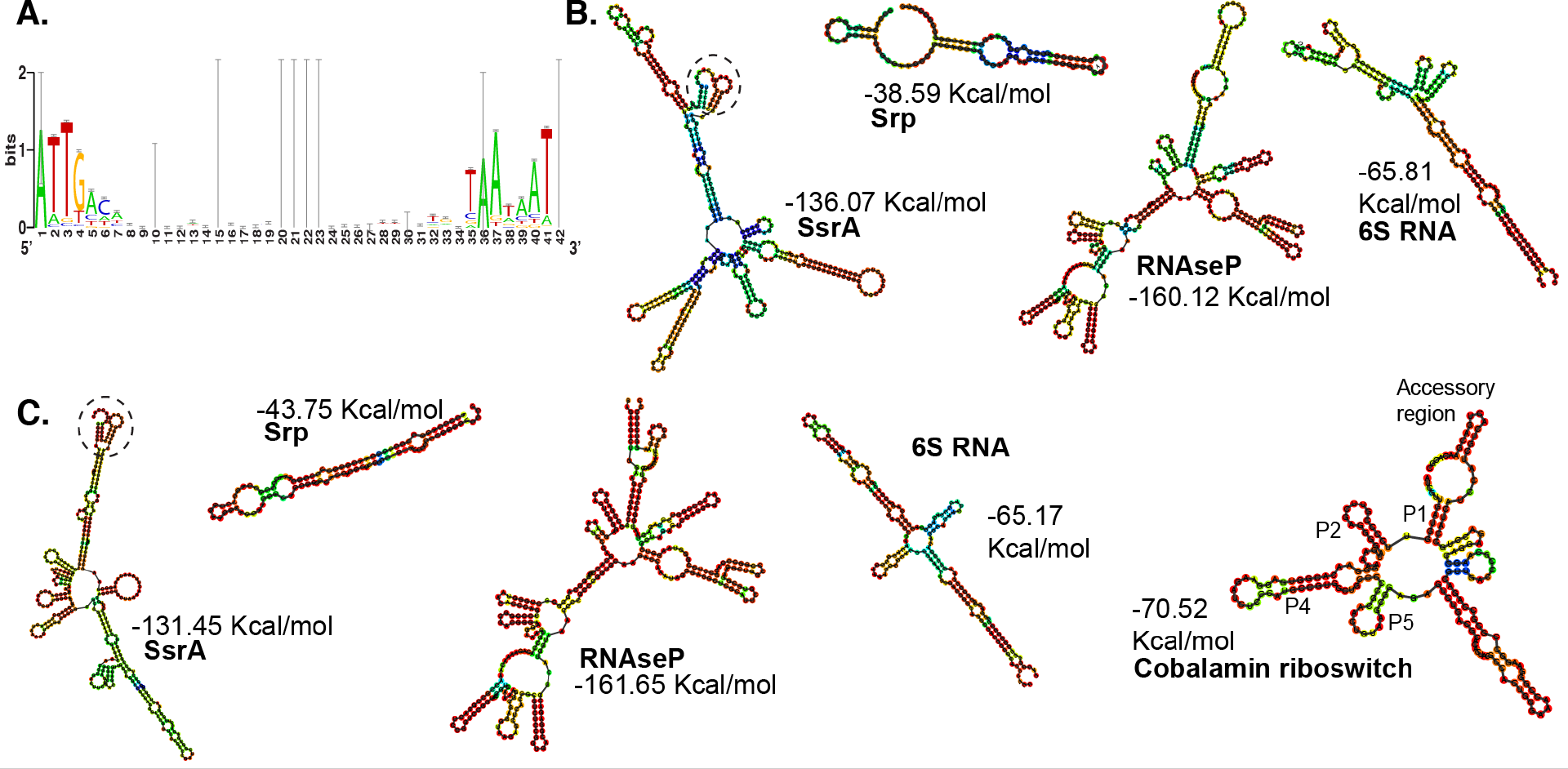
Predicted secondary structures of transcribed ncRNAs of known function in *Leptospirillum* sp. A) Sequence logo representation of the promoter structure in *Leptospirillum* group II UBA genotype. The alignment was constructed using the sequences of 207 predicted promoters. B) structures predicted in *Leptospirillum* group II UBA (Top structures). C) Structures predicted in *Leptospirillum* group III. Secondary structure prediction was done on the RNAfold webserver.

Transposases in *Leptospirillum* group II UBA and group III are among the most highly expressed genes (Supplementary Materials – R data). In addition, highly expressed multi-copy transposases are enriched in environmental than bioreactor samples (Supplementary Figure S2). Highly abundant transposase expression was reported in community proteomic analyses in acid mine drainage biofilms (26), and in community transcriptomic analyses (27, 28). It is possible that the movement of mobile elements is very important in natural acidophilic biofilms.

A few of the well-characterized ncRNAs in the Rfam database were identified in many genomes of acidophilic microorganisms, some of which are among the most highly expressed genes (Supplementary Table S3 and Figure 4). SsrA (aka transfer-messenger RNA or tmRNA) was detected in the *Leptospirilli* (Figure 4B-C), as well as in G-plasma, a plasmid, and in *Acidithiobacillus caldus,* the later representing only 0.003% of the community transcriptome in the sample in which it is most abundant (Adrift GS0, Supplementary Table S1). The RNAse P ncRNA was the most highly expressed ncRNA in the transcriptomic datasets, and was identified in most AMD organisms, including the fungus, the very low abundance *A. caldus* and *L. ferrooxidans* bacteria, and the archaeon C-plasma (Supplementary Table S3). The signal recognition particle (SRP) RNA was also highly expressed in the *Leptospirilli* (Figure 4B-C), *Ferroplasma* Type I and Type II, the fungus, and in G-, A-, E- and C-plasma. The 6S RNA (SsrS) was identified in the *Leptospirilli* (Figure 4B-C), the *Sulfobacillus*, and *A. caldus*. The cobalamin riboswitch was only identified in *Leptospirillum* group III and contains the expected conserved domains within the core region as described in other organisms (29) (Figure 4C). The cobalamin riboswitch has not yet been described in the genomes of any other member of the *Nitrospira* Phylum.

Among the ncRNAs associated to mobile elements detected in the transcriptomes are the Group II introns of *Leptospirillum* groups II and III, *Ferroplasma* Type II, and the Actinobacterial Bin 1 (Supplementary Table S3). In addition, the 5’ UTR region of several HNH-endonucleases in *Leptospirillum* groups II, III and IV, and *Ferroplasma* Type II contain the non-coding HEARO RNA, which, along with its associated endonuclease, constitute another type of mobile genetic element (30). The hgcC RNA, a non-coding RNA of unknown function, was detected in *Leptospirillum* group II UBA and group II 5way-CG, and in two viruses (Supplementary Table S3). Previously the hgcC RNA was reported only in Archaea (31), although it is also found in a few viruses in the Rfam database.

The small nucleolar RNA (snoRNA), a eukaryotic small RNA with few known homologs in Archaea (32) was detected in most Richmond Mine Archaea A-, C- and G-plasma, *Ferroplasma* Type I and Type II, and the fungus; while the crcB RNA (a fluoride riboswitch, (33)) was detected in the *Leptospirilli*, G-plasma, and in *Ferroplasma* Type II (Supplementary Table S3).

The TPP riboswitch binds thiamine pyrophosphate to regulate expression of thiamine biosynthesis and other related genes. These riboswitches have been identified upstream of multi-transmembrane hypothetical proteins in the genomes of *Thermoplasma acidophilum, Thermoplasma volcanicum, and Ferroplasma acidarmanus* (34). A-, G-, and E-plasma (members of the *Thermoplasmatales*), I-plasma and *Ferroplasma* Type II express the TPP riboswitch and reads within it have paired-reads to a downstream putative transporter likely regulating its expression (Supplementary Table S3).

### CRISPR Cas genes and loci expression occurs on both strands of DNA

Transcripts from CRISPR-associated Cas genes were detected in all samples for *Leptospirillum* groups II and III, the archaeon *Ferroplasma* Type I, and Archaea A- and G-plasma (Supplementary Table S4). Cas genes from *Leptospirillum* group II UBA and G-plasma showed higher expression in bioreactor samples, while those from *Leptospirillum* group III and group II 5way-CG showed higher expression in environmental biofilms (Figure 3E).

When surveying the transcriptomic reads using the CRASS software, transcripts containing CRISPR repeats from eight datasets were detected in *Leptospirillum* groups II and III, G-plasma, *Ferroplasma* Type I and Type II, the Actinobacterial bin, and plasmids (Supplementary Tables S4 and S5). Some additional CRISPR transcripts could not be assigned to an AMD genome based on known repeat sequences. The highest diversity of CRISPR repeats expressed was observed in the late growth-stage biofilm (4-way_GS2; Supplementary Table S4). This finding probably reflects the higher richness of late growth-stage biofilms as well as the activity of multiple closely related strains with slightly different CRISPR loci (and thus different phage/viral susceptibility). For *Leptospirillum* group II and G-plasma, the largest number of distinct spacer transcripts was detected in this same biofilm, consistent with a higher diversity of strains in the sample.

Transcriptomic reads from precursor CRISPR RNA (pre-crRNAs, as reviewed in (35)) aligned mostly at the trailer end (containing older CRISPR spacers) of the composite CRISPR loci from the assembled genome of *Leptospirillum* group II UBA (Figure 5A), and no repeats and spacer-carrying reads aligned to the CRISPR loci from the assembled *Leptospirillum* group III genome. These observations are likely due to the composite assemblies not capturing the dynamics of acquired spacers from closely related strains. For example, 407 of the 688 different spacer sequences recovered from *Leptospirillum* group II transcripts, and 5 of the 245 spacers detected in the transcriptome in *Leptospirillum* group III (Supplementary Table S5) had been recovered in a previous analysis of Richmond Mine CRISPR systems (36). Notably, while transcription of Cas genes occurs on the sense strand in which the genes were predicted, transcription of the CRISPR loci (repeat/spacer region) in *Leptospirillum* group II UBA occurs on the antisense strand (Figure 5A). Moreover, 4 of the 5 *Leptospirillum* group III CRISPR spacers and 404 of the 407 spacers in *Leptospirillum* group II UBA recovered here via strand-specific transcriptomics represent the reverse complement of the sequences previously reported in (36) (Supplementary Table S5).

**Figure 5.**
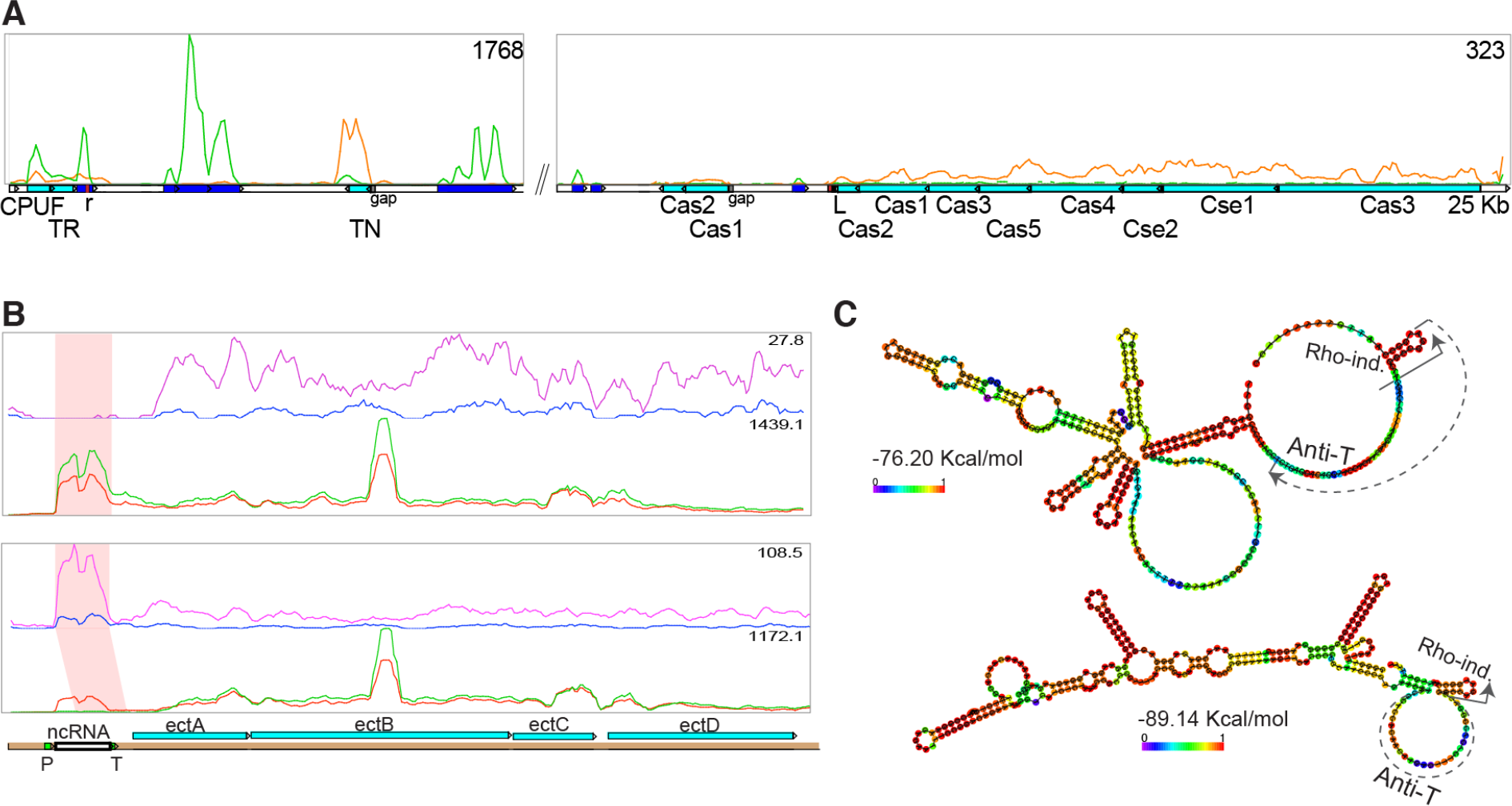
Expression of regulatory ncRNAs in *Leptospirillum* group II. A) Modified Artemis image showing the transcriptional expression of *Leptospirillum* group II UBA CRISPR loci. Expression of Cas genes occurs in the same orientation as the predicted genes on scaffold 8241 (orange line, (-) strand), while pre-crRNAs are *cis*-transcribed on the antisense strand (green line, (+) strand). CPUF: conserved hypothetical protein; TR: transcriptional regulator; TN: transposase. Pre-crRNAs are shown as blue boxes, predicted coding genes are represented by teal boxes, the CRISPR leader sequence (L) is represented by a black box downstream of Cas2 and is followed by the first repeat in the composite assembly (red box). The last repeat of the loci is also show as a red box next to the transcriptional regulator. The composite assembled CRISPR locus, including the Cas genes, is ~ 25 Kb in length. B-C) Novel ectoine riboswitch in *Leptospirillum* group II bacteria. The ectoine operon (teal blue arrows), and the riboswitch (ncRNA, white box) with its promoter (P) and Rho-independent terminator (T) are shown. The modified Artemis image in *Leptospirillum* group II UBA (B, top) and in *Leptospirillum* group II 5way-CG (B, bottom) shows the strand-specific transcriptional expression from two bioreactor samples (blue: R3_GS0; magenta: R3_GS1) and two environmental biofilms (red: AB10_GS0; green: AB10_GS1). The predicted secondary structure of the riboswitches are shown in: *Leptospirillum* group II UBA (C, top) and in *Leptospirillum* group II 5way CG (C, bottom) with their Rho-independent terminator (solid arrow) and an anti-terminator (dashed arrow).

### Expression of novel ncRNA

We observed transcription of long leader sequences upstream of many genes and operons in *Leptospirillum* groups II and III. One example is a transcribed 370 bp 5’ UTR of the ectoine biosynthesis operon in both *Leptospirillum* group II UBA and 5way-CG strains (Figure 5B-C). Ectoine is a compatible solute either synthesized or transported from the environment by many organisms during osmotic stress (37, 38). The length of the ncRNA upstream of the ectoine operon, the presence of paired reads from the ncRNA to the transcribed operon, and the presence of a putative promoter (TTGACA-N17-(A)A(A)A(C)T), a Rho-independent terminator (−6.30 Kcal/mol) and an anti-terminator (−7.03 Kcal/mol) suggest it is likely a riboswitch. The expression of the putative riboswitch in *Leptospirillum* group II UBA in two environmental biofilms is generally higher than that of the operon (Figure 5B, top panel, green and red curves). However, the transcript levels of the putative riboswitch in the bioreactor samples are much lower than those of the operon (Figure 5B, top panel, blue and magenta curves). These results suggest that the riboswitch may inhibit transcription of the operon in the environmental samples (preferred conditions for *Leptospirillum* group II UBA growth, see Figure 1) and enhance the ectoine operon transcription in bioreactor samples (where growth conditions are more stressful). The expression of the putative riboswitch in *Leptospirillum* group II 5way-CG shows a slight opposite trend than in *Leptospirillum* group II UBA (Figure 5B, bottom panel). The riboswitches of both bacteria share 80% identity at the nucleotide level and their predicted secondary structures look very different (Figure 5C).

Novel carbon monoxide dehydrogenase-associated ncRNAs (referred to here as CODH-ncRNA) were identified in long transcribed leader sequences in *Leptospirillum* group III and in *Leptospirillum* groups II UBA and 5way-CG genotypes (Figure 6 and Supplementary Figure S3). The CODH-ncRNA is expressed at similar abundance and contains paired-end reads that map to its downstream CO-dehydrogenase beta subunit gene (the only subunit identified in the *Leptospirilli*). Although terminators could not be predicted, the dip in the transcriptional expression of the CODH-ncRNAs before the ribosome binding site (RBS) of their associated CO-dehydrogenase (CODH) genes suggest that the CODH-ncRNA may be regulating CODH expression (Figure 6 and Supplementary Figure S3). Although the Archaeon *Ferroplasma* Type II has multiple copies of the full CODH operon and all subunits showed transcriptional expression, a CODH-ncRNA was not identified upstream of the operons (data not shown). Two of the three copies of the CODH-ncRNA in *Leptospirillum* group III and both copies in *Leptospirillum* group II are highly expressed in the late developmental-stage biofilms (Figure 6, left panel; Supplementary Figure S3A-D).

**Figure 6.**
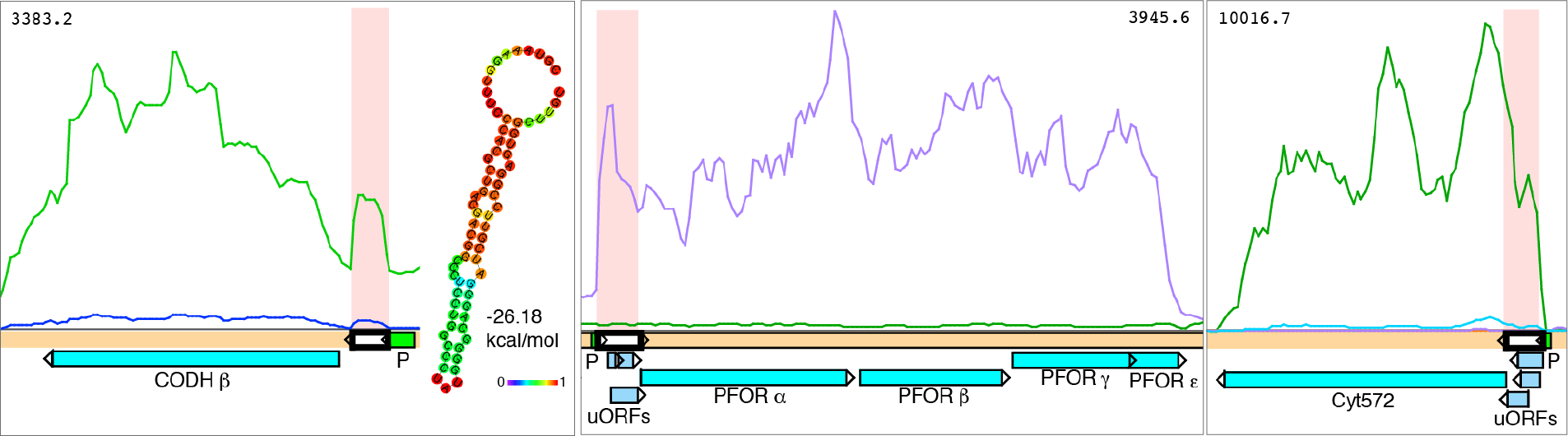
Expression of regulatory ncRNAs in *Leptospirillum* group III. Panels show the strand-specific expression of energy generation-related genes with their associated regulatory ncRNA. Coding genes are represented by teal arrows, ncRNAs by white arrows, putative uORFs by blue arrows, and promoters by green arrows. Left: CO dehydrogenase (CODH) beta subunit with its transcribed short (70 bp) ncRNA. The predicted secondary structure for the CODH-ncRNA is shown. Purple and green lines: R3_GS1 bioreactor biofilm; blue and teal lines: R3_GS0 bioreactor biofilm. Middle: pyruvate:ferredoxin oxidorectase (PFOR) operon with its transcribed 5’ UTR containing a putative promoter (P, green) and possible regulatory uORFs. Right: Cytochrome c 572 (Cyt572) with its transcribed 5’ UTR containing a putative promoter (P) and possible regulatory uORFs. The length of the putative regulatory uORFs ranges from 10 to 40 amino acids long.

We observed expression of many long leader sequences upstream of other important coding genes and operons in *Leptospirillum* groups II and III. For example, the transcribed 5’ UTR of the pyruvate ferredoxin oxidoreductase (PFOR) operon, cytochrome 572 and cytochrome 579 genes contain putative promoters and untranslated ORFs (uORFs) (Figure 6 and Supplementary Figure S3E-I). Other examples include expression of the 5’ UTR of the Uroporphyrin-III C-methyltransferase gene and of the Ribonuclease III gene in *Leptospirillum* group II UBA, with potential folded ncRNAs and predicted promoters (Supplementary Figure S3J-K).

## DISCUSSION

Community expression profiles inform about the ecology and physiology of actively growing organisms in their natural environment. Richmond Mine AMD biofilms develop in extremely acidic conditions and their characterization is vital to understanding the process of water acidification. Bioreactor conditions generally resemble environmental conditions from downstream AMD sites at higher pH and lower temperatures, as well as those from metal-release bioleaching systems. Therefore, analyses of environmental and laboratory-grown acidophilic biofilms improve our understanding of the dynamics occurring in acid drainage and bioleaching ecosystems.

Acid mine drain communities have been extensively studied for community membership, and proteomic and transcriptomic studies have been used to infer ecological roles of community members (reviewed in (20)). However, no reports have analyzed the transcriptional profiles of both coding genes and non-coding RNAs (ncRNAs) in acidophilic communities. Here, metatranscriptomic analysis of natural AMD biofilms and of acidophilic biofilms growing in laboratory bioreactors identified patterns in gene expression that correlate with organismal environmental preferences and reveal the potential roles of known and novel ncRNAs.

Differential gene expression analyses indicated that preferred growth conditions for *Leptospirillum* group II UBA and C75 genotypes, and the Archaeon G-plasma are environments at very low pH and high temperatures, while *Leptospirillum* group II 5way-CG and group III prefer higher pH and lower temperature conditions. AMD organisms appear to show base-level activity by expressing the whole genome, and *Leptospirillum*-associated viruses follow transcriptional expression profiles similar to that of their host. In addition, gradients of pH and temperature evident from differentially expressed genes, high expression of transposase genes, and the high diversity of CRISPR loci spacers suggest that AMD communities are highly dynamic ecosystems.

It has been suggested that Cas proteins and CRISPR loci primary transcripts are constitutively expressed, and their expression levels might be induced as invasion occurs (reviewed in (39)). Differential expression of CRISPR Cas genes in environmental vs. bioreactor biofilms point to some level of regulation, where Cas genes are up-regulated in less optimal environments for the organisms carrying them. *Leptospirillum* Group II Cas proteins, at the time considered hypothetical, were highly abundant in the first microbial community proteomic analysis (26) and expression of Cas genes was observed previously in *Leptospirillum* group III (18). When examining the strand-specific nature of the transcriptome, we observed that CRISPR loci (repeats and spacers) transcription occurs on the antisense strand while Cas genes are expressed on the sense strand in the *Leptospirilli*. Co-expression of sense and antisense mRNAs in plants was shown more effective at offering viral immunity than only sense or antisense expression of the relevant genes (40). Therefore, Cas genes and CRISPR loci expression on different strands may be used as a mechanism for effective viral immunity in the *Leptospirilli*. Although, differential expression of Cas genes and CRISPR loci on opposite strands has not yet been reported, bidirectionality of CRISPR loci transcription was observed in *Sulfolobus* sp., where the authors suggest the potential need for bidirectionality to neutralize the leader spacer RNAs in the absence of invading elements (41).

RNASeq analyses enable identification of regulatory ncRNAs, leader sequences and transcription start sites (TSS) for coding genes and operons. We confirmed expression of ncRNAs of known function in many high and low abundance organisms, including the RNAse P, SsrA and 6S ncRNAs previously predicted *in silico* in *Acidithiobacillus caldus* (42). In addition, some ncRNAs of unknown function and 5’ UTRs show expression levels similar to their neighboring genes and predicted operons, suggesting that expression of upstream non-coding regions might be regulating their neighboring genes. For example, we identified a novel riboswitch associated to the ectoine biosynthetic pathway in *Leptospirillum* group II. The riboswitch appears to inhibit transcription of the ectoine operon in favorable growth conditions while enhancing transcription during stress, suggesting that *Leptospirillum* group II synthesis of compatible solutes to tolerate unfavorable conditions is regulated by the ectoine riboswitch.

Other novel regulatory ncRNAs include a carbon monoxide dehydrogenase-associated ncRNA (CODH-ncRNA) in *Leptospirillum* group II and III. CO dehydrogenase, a nickel iron-sulfur protein, is usually part of a multi-protein complex which metabolizes carbon monoxide (CO) when the cells sense CO in the environment, as well as being used as electron and carbon sources by some bacteria (reviewed by (43)). The CODH beta-subunit, the only subunit identified in *Leptospirillum* groups II and III, has been reported as a single active subunit in metal-reducing *Geobacter* bacteria and in the genomes of two *Chlorobium* phototrophic bacteria (44). It is highly sensitive to oxygen, which might explain the high expression of both CODH and CODH-ncRNA transcripts in late developmental stage biofilms. It is possible that the CODH-ncRNA up-regulates expression of the CO dehydrogenase gene as biofilms mature.

Long transcribed 5’UTRs with putative upstream ORFs (uORFs) preceding energy-generation genes (e.g. PFOR and cytochrome c) likely regulate transcription and/or translation of their downstream genes and operons (Figure 6). Analysis of transcription start sites (TSS) in *Clostridium phytofermentans* have shown that the TSS of one *pfor* gene occurs 78 bp upstream of its start codon (45), a length similar to the putative TSS of the *pfor* operons in *Leptospirillum* sp. (Figure 6), while transcripts with long 5’ UTRs encoding putative ncRNAs and uORFs have been reported in *Haloferax volcanii* (46) and in *Mycobacterium* sp. (47). Overall, our results point to significant roles of regulatory ncRNAs in acidophilic communities and suggest that gene expression profiles appear to be a factor in ecological diversification.

## METHODS

Eight biofilms at different developmental stages (growth-stage (GS) 0, 0.5, 1, and 2) were collected from the A-, C-, AB-drift, and 4way locations within the Richmond Mine at Iron Mountain Mines, California (40°40′ 38.42″ N and 122″ 31′ 19.90″ W, elevation of approx. 900 m) (Table 1). AMD biofilm developmental stages were estimated visually based on thickness (23). In addition, biofilms were grown in the dark, at pH 1 and 37 °C in laboratory bioreactors, as described in (48) using inocula from within the A-drift location and mine drainage outflow (Table 1). Biofilms were harvested when reaching growth-stage 0, 0.5, and 1, snap-frozen in liquid nitrogen upon collection/harvest and stored at −80 °C.

Total RNA was extracted from frozen samples using two acid phenol-chloroform-isoamyl alcohol extractions and immediately purified using the RNEasy MinElute kit (Qiagen). Ribosomal RNA (rRNA) subtraction on 8 of the 13 biofilm samples was done using the MicrobExpress kit (Ambion). Good quality RNA (RIN > 7, assessed by a Bioanalyzer 2100 (Agilent Technologies)) from total RNA and rRNA-subtracted RNA were converted to cDNA as described by (49) in order to keep the strand-specificity of the transcriptome. Briefly, the technique involves adding deoxy-UTP in place of deoxy-TTP during synthesis of the second strand of cDNA. After Illumina library preparation, the second strand of cDNA is selectively digested allowing for sequencing of all molecules in the same direction, and sequences maintain the strand-specificity of the original RNA molecules (49). Resulting cDNA was fragmented using a Covaris S-system (Covaris, Inc.) to an average fragment size of 200 bp and sent to the University of California Davis for Illumina library preparation, digestion of the dUTP-containing strand, and sequencing. Five samples were sequenced using the GAIIx platform (75 bp, single end reads), while eight samples were sequenced using the HiSeq 2500 platform (100 bp, paired-end reads) (Table 1).

Low-quality bases were trimmed from the sequencing reads using the fastx_trimmer script (http://hannonlab.cshl.edu/fastx_toolkit/) and the sickle trimmer script with default parameters (https://github.com/najoshi/sickle) and reads < 40 bp in length were discarded. Trimmed reads were mapped to the SSU and LSU rRNA gene Silva database SSURef_102 (50) using Bowtie2 (51) with default parameters to separate ribosomal from non-ribosomal reads.

Non-ribosomal (non-rRNA) reads were mapped using Bowtie2 with default parameters to the available genomes of acidophilic bacteria: *Leptospirillum ferrooxidans* C2-3*, Leptospirillum* group II UBA type (*L. rubarum*), group II ‘5way-CG’ type, group II ‘C75’ type, group III (*L. ferrodiazotrophum*), group IV UBA BS, *Acidithiobacillus caldus* ATCC 51756, a *Sulfobacillus* bin, and two Actinobacterial bins (2, 3, 22, 52–56); archaea: A, C, D, E, G, and I-plasma, ARMAN-1, −2, −4, and −5, and *Ferroplasma* Type I and Type II (2, 4, 57–59); plasmids (53), a fungus (1); and nine viruses/phage (60). Mapped reads were then assembled into transcript fragments using the Cufflinks pipeline (61). The genome references for mapping are available at: http://genegrabber.berkeley.edu/amd/organisms.

The relative abundance of genomes and genomic fragments from AMD organisms in each sample was estimated as the coverage for all transcripts within a genome (read counts were normalized for genome length and total number of reads per sample) (Figure 1 and Supplementary Table S1). Assembled transcripts from all organisms were also searched vs. the Rfam database (62) for non-coding RNAs (ncRNA) of known function. Additionally, the transcriptional profiles of *Leptospirillum* groups II UBA, group II 5way-CG, group III, and of the archaea A-plasma, C-plasma, G-plasma, and *Ferroplasma* Type II, were visually inspected using Artemis (63). Transcribed regions that did not fall within a coding sequence were evaluated for the presence of non-annotated protein sequences using BlastX (64) vs. the non-redundant NCBI database. We scanned these regions for the presence of possible ribosome binding sites and start codons that could hint to hypothetical proteins not yet identified in the public databases. Transcribed intergenic regions that do not appear to encode for protein sequences, based on the above criteria, were labeled as potential non-coding RNA (ncRNA). Bacterial promoters were predicted upstream of manually evaluated ncRNAs using BPROM (65) and rho-independent terminators were evaluated with ARNOLD (66). Riboswitch motifs were predicted using the RibEx webserver (67), and secondary structure prediction of the ncRNAs was done using the RNAfold webserver (68).

Correlation analyses of assembled transcript abundances of non-rRNA reads that mapped to the predicted genes of AMD organisms were done on samples for which total and rRNA-depleted reads were obtained (Supplementary Figure S4). Transcript abundances correlated well for most samples (R^2^ values range from 0.74 to 0.95) hence, non-rRNA reads from rRNA-depleted RNA were pooled with those non-rRNA reads from their corresponding total RNA. The correlation between rRNA-depleted and total RNA transcripts from an early developmental stage bioreactor sample, R1_GS0, was low (and dispersion of points was high), likely due to the much shorter fragments assembled from the total RNA sample (Supplementary Figure S4H). Given that the correlation between transcript abundances was slightly positive, and that transcript length improved in the rRNA-depleted sample, reads from total and rRNA-depleted RNA were also pooled for R1_GS0.

Tables of AMD genes read counts were used for differential expression analyses in R (69). Read counts were normalized for gene length using the function “withinLaneNormalization” from the EDASeq package (70). NMDS ordination was performed on Bray-Curtis distance matrices of variance-stabilizing transformed gene abundance tables with the formula: genes ~ sequencing type + environment type, using the DESeq2 R package (71).

Putative RpoD promoters were predicted upstream of expressed genes in *Leptospirillum* group II UBA genotype by manual inspection of the genome using Artemis (63) along with the visualized transcriptome data. Predicted promoters were listed if they were less than 50 bp upstream of the 5’ end of the mapped reads of a transcript and matched 8 out of 12 bp of the standard *E. coli / B. subtilis* RpoD motif (TTGACA_N_16-19__TATAAT).

Hierarchical clustering of abundances from predicted genes was done using the software Cluster 3.0 for Mac OSX, centering genes and samples by the median, using the Spearman Rank Correlation similarity matrix, and Average Linkage as clustering method (72). Clusters and heat maps were visualized using the Java TreeView software (73). Gene trees for *Leptospirillum* spp. transposases were constructed using the MABL website (74). CRISPR loci from transcriptomics reads were reconstructed using the CRASS program (75).

## DATA AVAILABILITY

Raw sequencing reads were submitted to the NCBI sequence reads archive (SRA) under accession number SRP026490 (76). R code and data tables are available at https://github.com/dgoltsman/acid-mine-drainage.

## ACKNOWLEDGEMENTS

We thank the late TW Arman, President of Iron Mountain Mines Inc., Mr R Sugarek (US Environmental Protection Agency) for site access, and Mr R Carver for on-site assistance. This project was funded by the U.S. Department of Energy, through the Carbon-Cycling (DE-FG02-10ER64996) program.

## SUPPLEMENTARY MATERIALS LEGENDS

1. Supplementary Table S1. Genome-wide relative abundance of transcripts from AMD genomes.

2. Supplementary Table S2. Predicted promoters upstream of transcribed genes in *Leptospirillum* group II UBA genotype.

3. Supplementary Table S3. Transcripts containing ncRNAs with predicted functions.

4. Supplementary Table S4. Summary of CRISPR spacers recovered from paired-end, strand-specific transcriptomics datasets.

5. Supplementary Table S5. CRISPR CRASS output table and spacer matches to previous CRISPR analyses.

6. Supplementary Figure S1. Differential expression analysis of genes from AMD genomes.

7. Supplementary Figure S2. Gene tree of transposases expressed in *Leptospirillum* sp.

8. Supplementary Figure S3. Examples of expression of leadered transcripts in *Leptospirillum* sp.

9. Supplementary Figure S4. Transcript abundance plots from total vs. rRNA-depleted RNA samples.

